# Persistent Neurological Deficits in Mouse PASC Reveal Antiviral Drug Limitations

**DOI:** 10.1101/2024.06.02.596989

**Authors:** Abhishek Kumar Verma, Shea Lowery, Li-Chin Lin, Eazhisaivallabi Duraisami, Juan E. Abrahante Lloréns, Qiang Qiu, Marco Hefti, C. Ron Yu, Mark W. Albers, Stanley Perlman

**Affiliations:** Department of Microbiology and Immunology, University of Iowa, Iowa City, IA 52242; Iowa Neuroscience Institute, University of Iowa, IA, USA 52242; Department of Neurology, University of Iowa, Iowa City, IA 52242; Minnesota Supercomputing Institute, University of Minnesota, Minneapolis, MN; Stowers Institute for Medical Research, Kansas City, MO 64110; Department of Pathology, University of Iowa, Iowa City, IA 52242; Department of Neurology, Massachusetts General Hospital, Harvard Medical School, Boston, MA

**Keywords:** SARS-CoV-2, Brain, Anosmia, Tyrosine Hydroxylase, Substantia Nigra, Neurodegeneration, Olfactory Bulb, Microglia, Inflammation

## Abstract

Post-Acute Sequelae of COVID-19 (PASC) encompasses persistent neurological symptoms, including olfactory and autonomic dysfunction. Here, we report chronic neurological dysfunction in mice infected with a virulent mouse-adapted SARS-CoV-2 that does not infect the brain. Long after recovery from nasal infection, we observed loss of tyrosine hydroxylase (TH) expression in olfactory bulb glomeruli and neurotransmitter levels in the substantia nigra (SN) persisted. Vulnerability of dopaminergic neurons in these brain areas was accompanied by increased levels of proinflammatory cytokines and neurobehavioral changes. RNAseq analysis unveiled persistent microglia activation, as found in human neurodegenerative diseases. Early treatment with antivirals (nirmatrelvir and molnupiravir) reduced virus titers and lung inflammation but failed to prevent neurological abnormalities, as observed in patients. Together these results show that chronic deficiencies in neuronal function in SARS-CoV-2-infected mice are not directly linked to ongoing olfactory epithelium dysfunction. Rather, they bear similarity with neurodegenerative disease, the vulnerability of which is exacerbated by chronic inflammation.

## Introduction

The global outbreak of COVID-19, caused by the severe acute respiratory syndrome-coronavirus-2 (SARS-CoV-2), has resulted in infection of over 773 million people and 7 million deaths worldwide reported to the World Health Organization (https://covid19.who.int/), as of January 4, 2024. While COVID-19 primarily involves the respiratory system^1,2^, several studies indicate that the central nervous system (CNS) is affected during acute and chronic SARS-CoV-2 infection, with consequent neurological and psychiatric complications^3–6^. Commonly reported manifestations include cognitive dysfunction, headache, loss and/or distortion of smell and taste, encephalopathy, delirium, strokes, seizures, neuropathy, and myopathy^7–14^. Less frequent problems include abnormal movements, psychomotor agitation, syncope, and autonomic dysfunction^15–17^. Many of these symptoms/signs resolve during convalescence, but some symptoms persist for extended periods of time, such as cognitive disturbances, neuropsychiatric symptoms, fatigue, insomnia, headache, loss of memory and anosmia/ageusia^18–21^. Importantly, the underlying biological mechanisms responsible for these persistent abnormalities remain unknown.

Systemic or localized inflammation has been implicated in SARS-CoV-2 neurological disease. For example, analysis of cerebrospinal fluid from acutely infected patients with neurological symptoms revealed increased levels of various cytokines^22–24^. SARS-CoV-2-induced neurological disorders does not seem to be a direct encephalitis since most reports demonstrate that the virus does not directly invade the central nervous system (CNS). It is possible that elevated levels of proinflammatory mediators in the blood can signal to the brain via hematogenous and neural pathways, i.e., peripheral inflammation may initiate neuroinflammatory events to affect brain function and contribute to SARS-CoV-2-associated neurological changes^25^. This possibility has not been thoroughly tested. Patients with Parkinson’s disease experience worsening of symptoms after SARS-CoV-2 infection^26–29^.

In patients and experimental animals, SARS-CoV-2 infect sustentacular cells in the olfactory epithelium (OE), which provide support for olfactory sensory neurons (OSN)^30–32^. Their dysfunction may cause anosmia through inflammatory responses^33,34^. Additionally, patients with previous SARS-CoV-2 infections manifest decreased global brain size and grey matter thickness, particularly in areas connected to the primary olfactory cortex on MRI scans after the acute phase of the infection^35,36^. SARS-CoV-2 infection may impact OB function through damage to the OE, but this link has not been clearly established.

Indeed, questions remain as to whether peripheral dysfunction can directly cause decreased central CNS function. In patients and experimentally infected animals, brain areas not considered directly connected to the olfactory system are affected^35,37,38^. Some of these sites, including the substantia nigra (SN), are affected in neurodegenerative disease, such as Parkinson’s Disease (PD)^39,40^. These observations raise the possibility that these brain areas may be vulnerable to pathogenesis independent of virus infection. That is, viral infection impinges on pre-existing vulnerability to exacerbate disease progress. It is plausible that certain cell types, e.g., dopaminergic neurons, are more susceptible to inflammatory responses. Consistent with this notion, the OB, which is rich in resident dopaminergic neurons, is affected in PD^41,42^. Chronic inflammation in the OB of PD patients can lead to olfactory dysfunction prior to the development of other symptoms^41–44^. A role for entry of environmental toxins or pathogens via the olfactory mucosa has been proposed as a contributory factor to both PD and Alzheimer’s Disease (AD)^45^. Similarly, there was an increased risk of PD after infection with another virus, the influenza virus that caused the 1918 pandemic^46–48^. In these cases, virus infection may cause brain dysfunction independent of anosmia. However, little is known about whether neurological dysfunction occurs and persists if the initial insult in the olfactory mucosa resolves. Thus, it is critical to understand whether there is an increased risk of neurodegenerative disease in COVID-19 survivors and whether it is related to persistent anosmia.

Here, we infected mice with a well-characterized mouse-adapted SARS-CoV-2 (SARS2-N501Y_MA30,_ referred to as SARS-CoV-2 herein) that infects the respiratory tract but not the CNS^49^. We focused on neurotransmitter and inflammatory molecule expression in the OB and SN since these sites are commonly involved in neurodegenerative disease and correlated these changes with behavioral changes and microglia gene expression. The results indicate that SARS-CoV-2 infection results in long term effects that are analogous to changes observed in patients with neurodegenerative disease and that these changes occur in the presence of treatment with anti-viral agents. We also show that changes in neurotransmitter expression in infected mice were present in the brains of deceased COVID-19 patients.

## Results

### Alterations in OB gene expression at late times after infection

We previously showed that OSN function in mice was impaired after acute SARS-CoV-2 infection^30^. OSN activity is correlated with the expression of tyrosine hydroxylase in the OB^50,51^. As a consequence of OE infection, we found that tyrosine hydroxylase (TH) expression was decreased in the OB during the acute phase^30^. To assess whether TH levels in the OB were chronically diminished by SARS-CoV-2 infection, we infected 14–18-week-old C57BL/6 mice with a sublethal dose of SARS-CoV-2 or PBS. Infected mice developed mild disease with 10-15% weight loss and 90% survival (Extended data Figure 1a). Brains were harvested at 120 days post-infection (dpi) and numbers of TH+ cells in the OB of mice were determined by immunostaining (Figure 1a). We observed a significant decrease in the total numbers of TH-positive cells in the glomeruli of the OB compared to mock-infected mice (Figure 1a). The decrease in TH was further confirmed by quantitative PCR analysis of whole OB for TH gene expression at 30 dpi and 120 dpi (Extended data Figure 1c, Figure 1b). Since inflammation has been implicated in diminished TH expression in other conditions, we next analyzed OB for the expression of proinflammatory cytokines. The data showed significant upregulation of IFN-β, IL-6 and TNF at 30 dpi whereas IFN-β, MDA-5, and NLRP3, were upregulated at 120 dpi. We also confirmed the complete absence of SARS-CoV-2 RNA by measuring N-gene expression in the OB (Extended data Figure 1b). Furthermore, as inflammation in the brain is associated with microglia activation, we investigated the activation of microglia through Iba-1 staining. We observed increased microglia/macrophage numbers in the OB at 120 dpi compared to mock-infected mice (Figure 1d) and a trend towards increased numbers of activated microglia (Extended data Figure 1d). Subsequently, we examined the numbers of microglia branches and junctions in mock and 120 dpi samples using skeletonize plugins in ImageJ and found an increased number of microglia with at least three junctions at 120 dpi (Extended data Figure 1e). However, there was no discernible change in the overall count of branches between the two sets of samples (Extended data Figure 1f). Further investigation found a reduction in branch length in the infected samples, providing additional evidence for microglial activation. Together, these data demonstrate that SARS-CoV-2 infection induces persistent OB alterations, including decreased TH expression, elevated cytokine levels, and increased numbers and activation of microglia, suggesting long-term consequences on olfactory function and neural inflammation. Changes in olfaction are often an early sign of neurodegenerative diseases such as Parkinson’s disease (PD) and Alzheimer’s disease (AD)^41,42,44^, so we next probed gene expression in the substantia nigra (SN), a major site impacted by PD^52–56^ using the same approach as described for the OB. SN is rich in dopaminergic neurons, which are decreased in PD.

**Figure 1.**
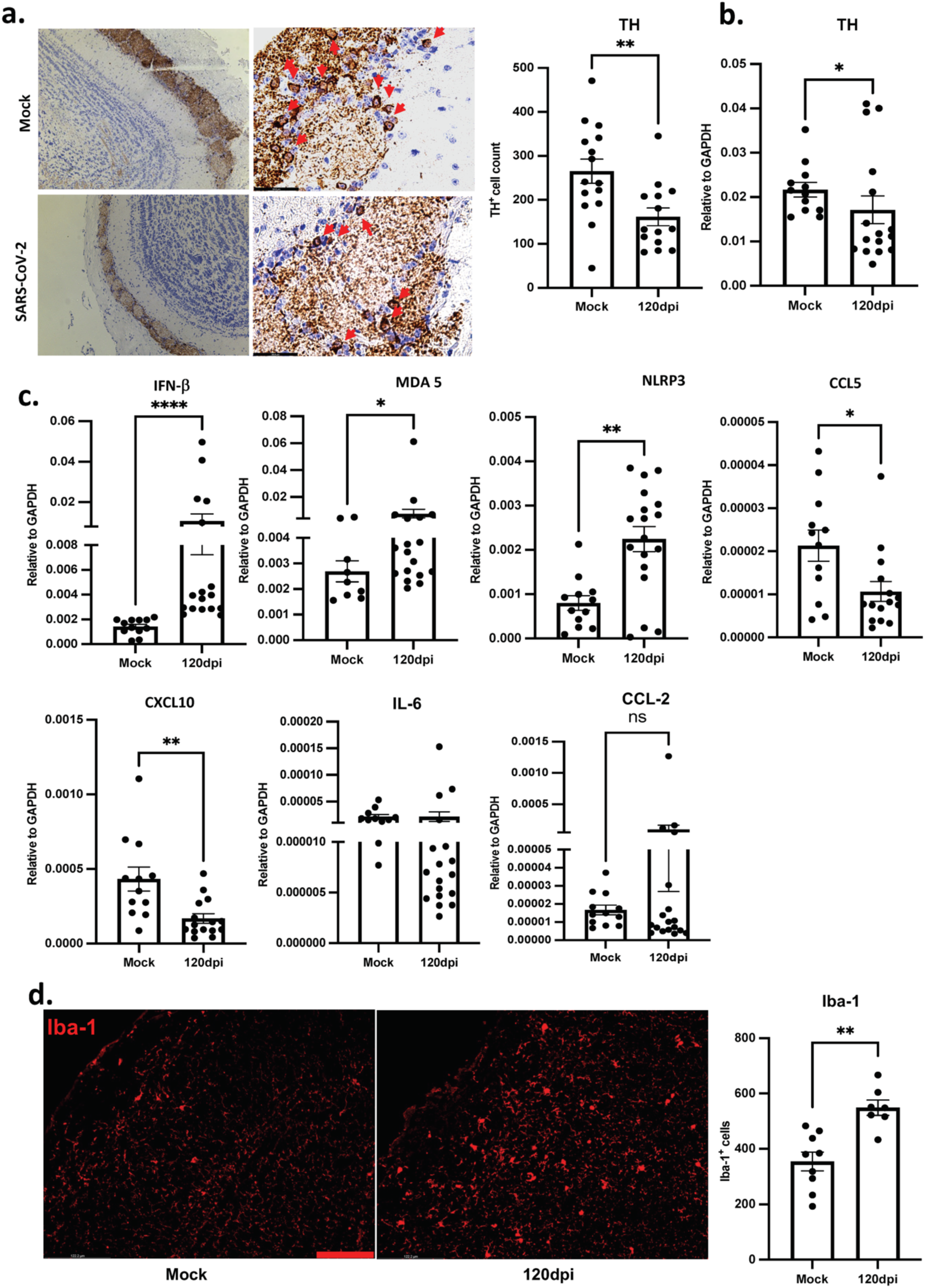
Loss of TH-positive cells and increased neuroinflammation in the OB at 120 dpi. (a) 4–5-month-old C57BL/6N mice were infected intranasally with 1000pfu SARS-COV-2. (left) Comparatively localized medial OB sections from infected and mock-infected animals were stained for TH. (right) Summary data of numbers of TH+ cells in periglomerular cells. Data represent mean ± SEM of results pooled from 3 independent experiments: mock (15 mice) and 120 dpi (14 mice). Data were analyzed using a Mann-Whitney U-test, \*\**P* < 0.01. Scale bar: 50 μm (b) OB mRNA was analyzed for TH expression by qPCR. Data represent mean ± SEM of results pooled from 3 independent experiments: mock (12 mice) and 120 dpi (16 mice). Data were analyzed using a Mann-Whitney U-test, \**P* < 0.05. (c) Proinflammatory cytokine mRNA expression was analyzed using qPCR. Data represent mean ± SEM of results pooled from 2 independent experiments. mock (10-12 mice) and 120 dpi (15 mice). Data were analyzed using a Mann-Whitney U-test. \**P* < 0.05, \*\**P* < 0.01, \*\*\*\**P* < 0.0001. (d) Myeloid cells were stained for Iba1 (red). Three to six fields from mock (8 mice) and 120 dpi (7 mice) were analyzed. Left. Representative sections from control and infected mice at 120 dpi are shown. Right. Summary data show numbers of Iba1^+^ cell in the OB. Data are mean ± SEM of results pooled from 2 independent experiments with 3 mice per group. Data were analyzed using a Mann-Whitney U-test. \*\**P* < 0.01. Scale bar: 50 μm.

### Alteration in gene profile in substantia nigra

We first confirmed the absence of SARS-CoV-2 in the SN by RT-PCR (Extended data Figure 1f). We then assessed the number of TH+ cells at this site and observed a significant decrease in the number of TH+ cells in infected compared to control SN (Figure 2a). The decrease in TH was confirmed using quantitative PCR analysis, which showed a decrease in TH mRNA in the SN at 120 dpi compared to mock-infected mice (Figure 2b).

**Figure 2.**
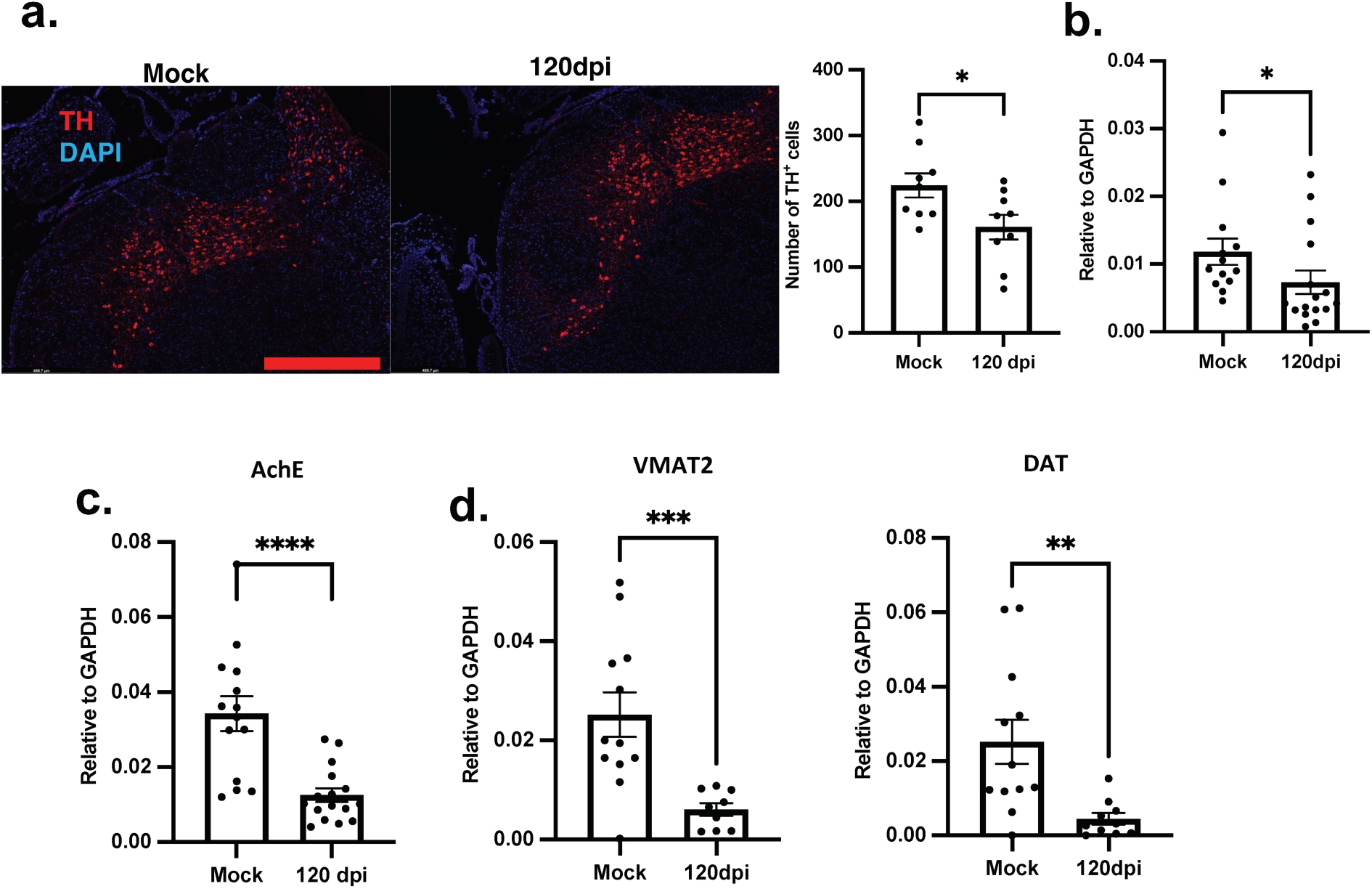
Loss of TH+ cells and changes of associated genes in substantia nigra. (a) Sections from control and infected brains were prepared at 120 dpi and control mice and stained for TH expression. Numbers of TH+ cells in SN of mice were quantified as described in Materials and Methods. Summary data show numbers of TH+ cells in the SN. Data represent mean ± SEM of results pooled from 2 independent experiments: mock (9 mice) and 120 dpi (9 mice). Data were analyzed using a Mann-Whitney U-test, \**P* < 0.05. Scale bar represents 490µm (b) TH mRNA expression in the SN was analyzed using qPCR. Data show mean ± SEM of results pooled from 3 independent experiments: mock (12 mice) and 120 dpi (15 mice). Data were analyzed using a Mann-Whitney U-test, \**P* < 0.05. (c) RNA was prepared from the SN of infected and uninfected mice as described in Materials and Methods. AchE mRNA expression was analyzed using qPCR. Data represent mean ± SEM of results pooled from 3 independent experiments: mock (12 mice) and 120 dpi (13 mice). Data were analyzed using a Mann-Whitney U-test, \**P* < 0.05. (d) DAT2 and VMAT mRNA expression in the SN was analyzed by qPCR. Data represent mean ± SEM of results pooled from 2 independent experiments: mock (11 mice) and 120 dpi (9 mice). Data were analyzed using a Mann-Whitney U-test, \**P* < 0.05.

Second, imbalances in acetylcholine levels are known to play a critical role in neurodegenerative diseases, with AD or PD patients having low levels of acetylcholine in the brain^57,58^. To determine whether cholinergic neurons were also affected in the SN, we performed qPCR for acetylcholinesterase (AChE) gene expression and found significantly lower levels in infected compared to control mice (Figure 2c).

To determine whether these effects on neurotransmitter expression were specific to the SN or were generalized, mRNA levels of TH and choline acetyltransferase (ChAT) were assessed in the brain after the OB, SN, and cerebellum were removed. We detected no significant differences in TH, AChE and ChAT mRNA levels in these samples when brains harvested at 120 dpi or from mock-infected mice were compared (Extended data Figure 1g and 1h). Since these results indicated that neurotransmitter expression in the SN was specifically affected by SARS-CoV-2 infection, we next measured levels of several mRNAs associated with dopamine expression, including dopamine transporter (DAT) and vesicular monoamine transporter 2 (VMAT2), as these proteins are decreased in PD. The results showed diminished DAT and VMAT2 mRNA expression in infected compared to control samples (Figure 2d).

Dopaminergic neurons of the substantia nigra are particularly vulnerable to neuroinflammation^53–56^. As neurotransmitter changes were observed in the SN, we next investigated inflammatory gene expression in the SN obtained from control and infected mice at 30 dpi and 120 dpi. Significant increases in IL-6 and MDA 5 mRNA were observed at 30 dpi whereas significant increases were observed in IFN-β and MDA5 levels at 120 dpi (Figure 3a). Given that a decrease in TH expression in the substantia nigra has been linked to dysregulation of several functional groups of genes involved in the PD^59^, we assessed expression of genes such as PTEN induced putative kinase 1 (PINK), Parkinson disease protein 7 (PARK7), ubiquitin C-terminal hydrolase L1 (UCHL), Leucine-rich repeat kinase 2 (LRRK2) and synuclein alpha (SNCA), which are associated with PD development. We found a significant decrease in PINK, PARK7, UCHL, and SNCA, but not in LRRK2 mRNA at 30 dpi and 120 dpi (Figure 3b).

**Figure 3.**
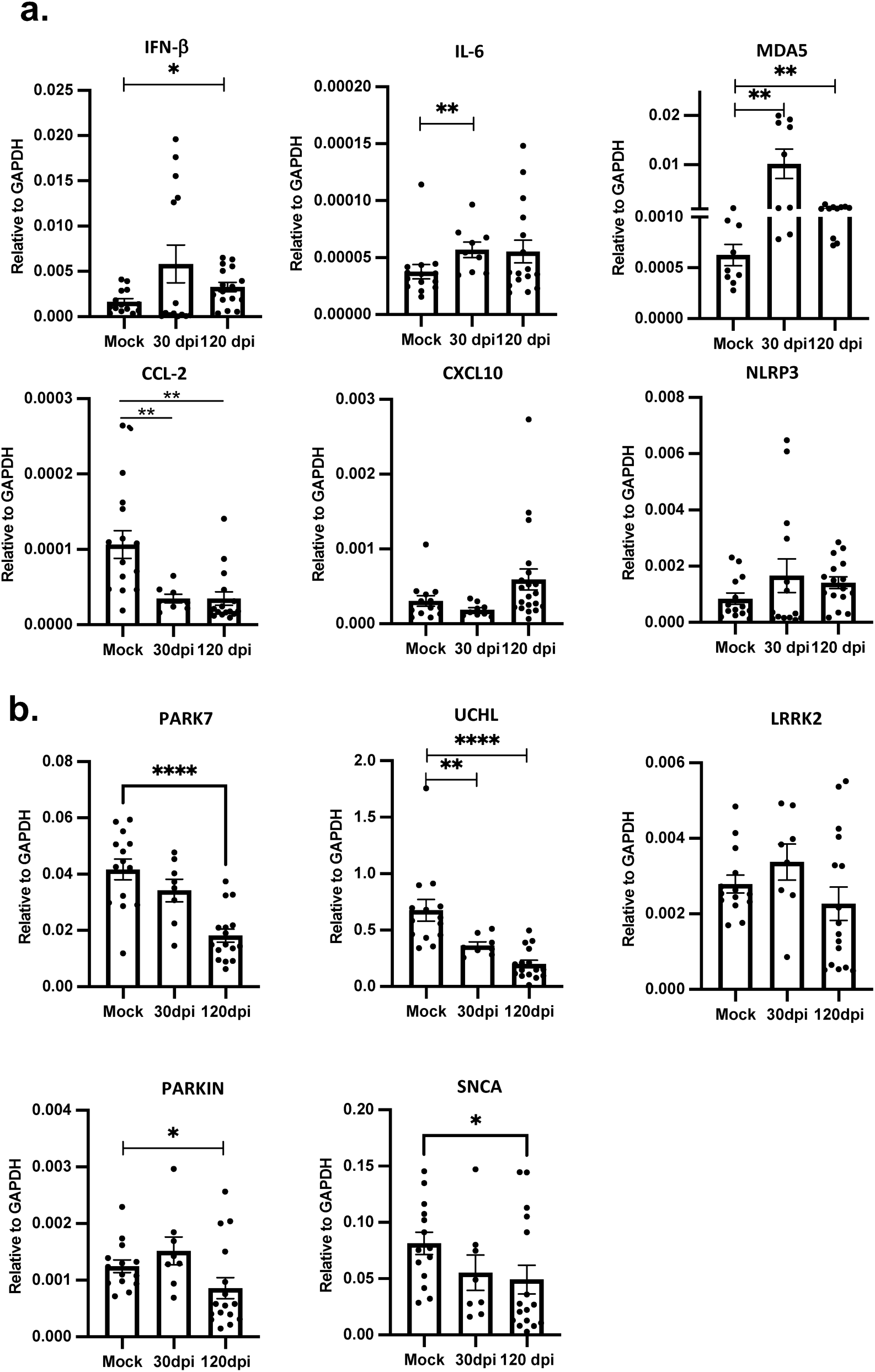
Neuroinflammation in the SN. (a) RNA was prepared from SN isolated from infected (30dpi and 120 dpi) and uninfected mice as described in Materials and Methods and analyzed for proinflammatory cytokine mRNA expression by qPCR. Data represent mean ± SEM pooled from 2 independent experiments: mock (10-12 mice), 30 dpi (8 mice) and 120 dpi (14 mice). Data were analyzed using a Mann-Whitney U-tests. \**P* < 0.05, \*\**P* < 0.01. (b) mRNA expression for genes associated with Parkinson’s Disease was analyzed using qPCR. Data represent mean ± SEM of results pooled from 2 independent experiments. mock (10-12 mice), 30 dpi (8 mice) and 120 dpi (14 mice). Data were analyzed using a Mann-Whitney U-test. \**P* < 0.05, \*\**P* < 0.01 \*\*\*\**P* < 0.0001.

### RNAseq analyses of myeloid cells

These targeted approaches of gene expression indicate that, despite the absence of infectious virus in the brain, SARS-CoV-2 infection has long term effects on gene expression in the brain. To obtain a broader view of alterations in expression, we next focused on total brain CD11b+ myeloid cells which are implicated in abnormal responses in the SARS-CoV-2 infected CNS in patients^22,60^. At 100 dpi, notable changes in gene expression in CD11b+ cells were observed. Expression of 99 genes was upregulated and of 147 genes was downregulated compared to mock-infected samples (Figure 4a). Figure 4b displays a heatmap depicting differentially expressed genes associated with inflammation. The upregulation of proinflammatory molecules such as TNF, CCL2, and CXCL10 was consistent with a persistent inflammatory state in the brain, present even three months post-infection. Next, we performed canonical pathway analysis using Ingenuity Pathway Analysis (IPA) at –2.5 to +2.5-fold change. Our data showed that the most highly activated pathway was the “Pathogen-induced cytokine storm signaling pathway”, suggesting a robust inflammatory response driven by CD11b cells (Figure 4c). Furthermore, disease and function analysis revealed the activation of inflammatory response and demyelinating pathways (Extended data Figure 2a), which is unexpected given that the brain was not directly infected with SARS-CoV-2. Kyoto Encyclopedia of Genes and Genomes (KEGG) pathway enrichment analysis identified the upregulation of genes associated with coronavirus disease. Together, these analyses point to an impact of systemic inflammation on brain function.

**Figure 4.**
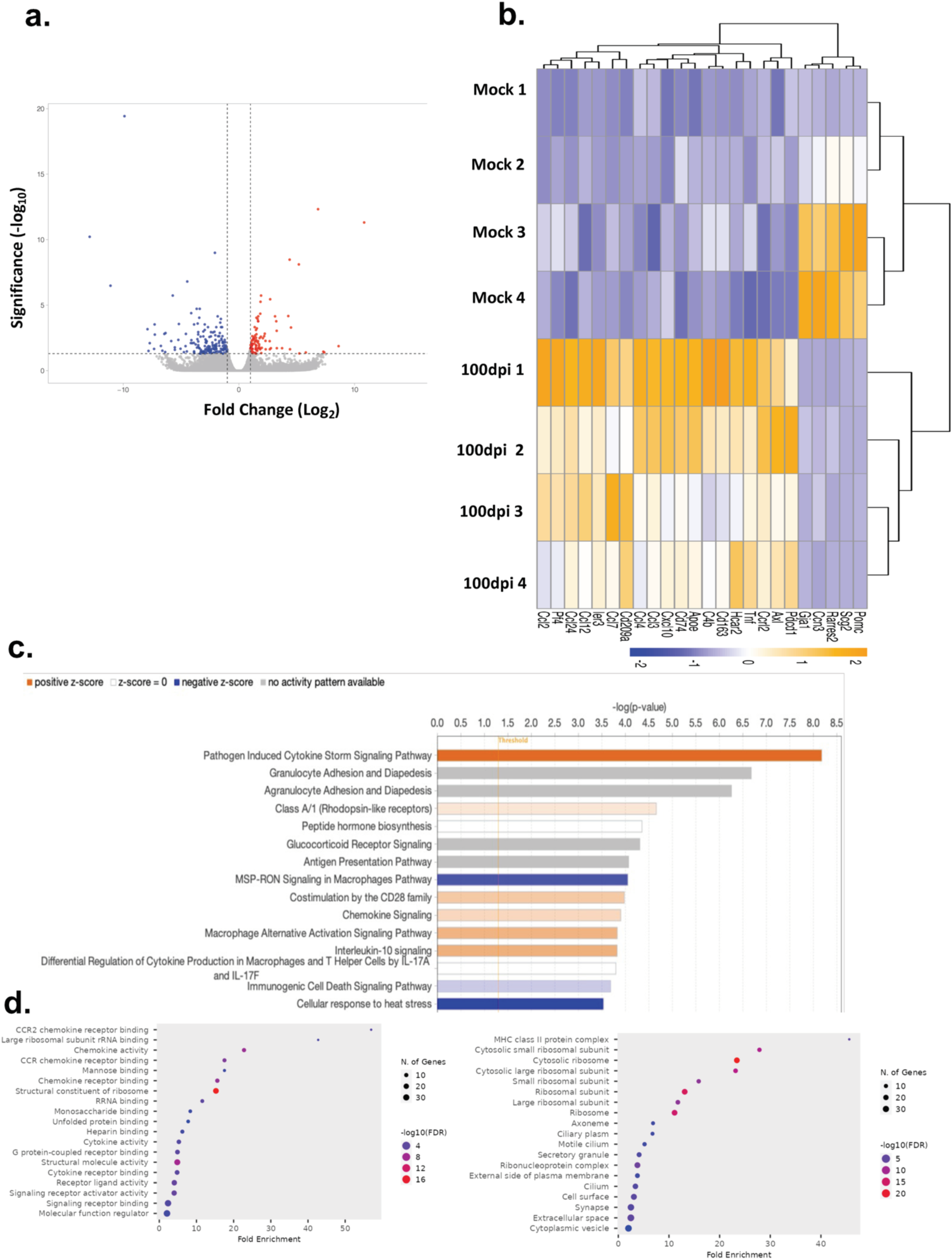
Differential gene expression in CD11b+ cells from SARS-COV-2 and mock-infected brains. CD11b+ cells were prepared from the brains of uninfected and infected mice at 100 dpi. RNA was prepared and analyzed by RNAseq as described in Materials and Methods. (a) Volcano plot depicting 246 differentially expressed genes in the CD11b+ cells of SARS-COV-2 infected mice in comparison with mock mice. Adjusted p-value ≤ .05, |fold-change| ≥ 2. (b) Heat map of differentially expressed inflammation-associated genes in SARS-COV-2–infected versus mock-infected CD11b+ samples. The scaled expression value (row Z score) is shown in a blue-red color scheme with red indicating higher expression, and blue lower expression. (c) Genes expressed at significantly higher levels in the CD11b+ cells were significantly enriched in Canonical Pathway Gene sets. X-axis denotes statistical significance as measured by negative logarithm of p-value. The ratio of differentially expressed genes was analyzed for statistical significance using Benjamini-Hochberg-corrected p-values < 0.05, |fold-change| ≥ 2.5. The red and blue bars represent categories for which specific functions are activated or repressed, respectively. (d-e) Gene ontology enrichment analysis for “Cellular Components” and “Molecular Functions pathway in CD11b+ cells are shown. Enrichment *p* values (corrected using the weighted Fisher’s method) <.05 are shown. The size and the color of each dot are proportional to the number of differentially expressed genes and the p-value respectively.

In addition, RNAseq analysis revealed upregulation of antigen processing and presentation pathways in CD11b cells, accompanied by significantly increased MHC-II expression (Figure 4d, right panel). Molecular function analysis identified upregulation of CCR2 receptor binding and chemokine receptor activation pathways, suggestive of enhanced immune cell migration and sustained inflammation in the brain (Figure 4d, left panel). We validated these results by RT-qPCR using CD11b+ cell RNA isolated from mouse brains at 100 dpi. These analyses confirmed increased expression of CCL2, CXCL10, TNF, and IFN-β ( Extended data Figure 2b) and are consistent with results from the RNAseq analyses.

### SARS-CoV-2 induces behavioral alterations in mice at 120 dpi

To determine whether the observed changes in gene expression in the SN effected changes in mouse motor function, we performed behavioral testing. Mice were subjected to rotarod testing, in which they were placed on a rod that was rotated at increasing speeds. The maximum speed prior to falling off the rod was recorded. We found a significant decrease in the speed achieved by infected mice before falling, as compared to mock-infected mice (Figure 5a). As another approach to assessing motor behavior, we performed open field testing (OFT) using control mice and mice at 40 and 100 dpi. The OFT provides qualitative and quantitative measurements of exploratory and locomotor activity in rodents^61^. Our results showed a decrease in the total distance covered by the mice at both 40 dpi and 100 dpi, indicating altered motor function (Figure 5b). We also observed that infected mice avoided the center part of the apparatus at 40 dpi but not 100 dpi, suggestive of some degree of increased anxiety (Figure 5b). These findings suggest that SARS-CoV-2 infection leads to alterations in the SN that affect normal mouse behavior, including motor and affective function.

**Figure 5.**
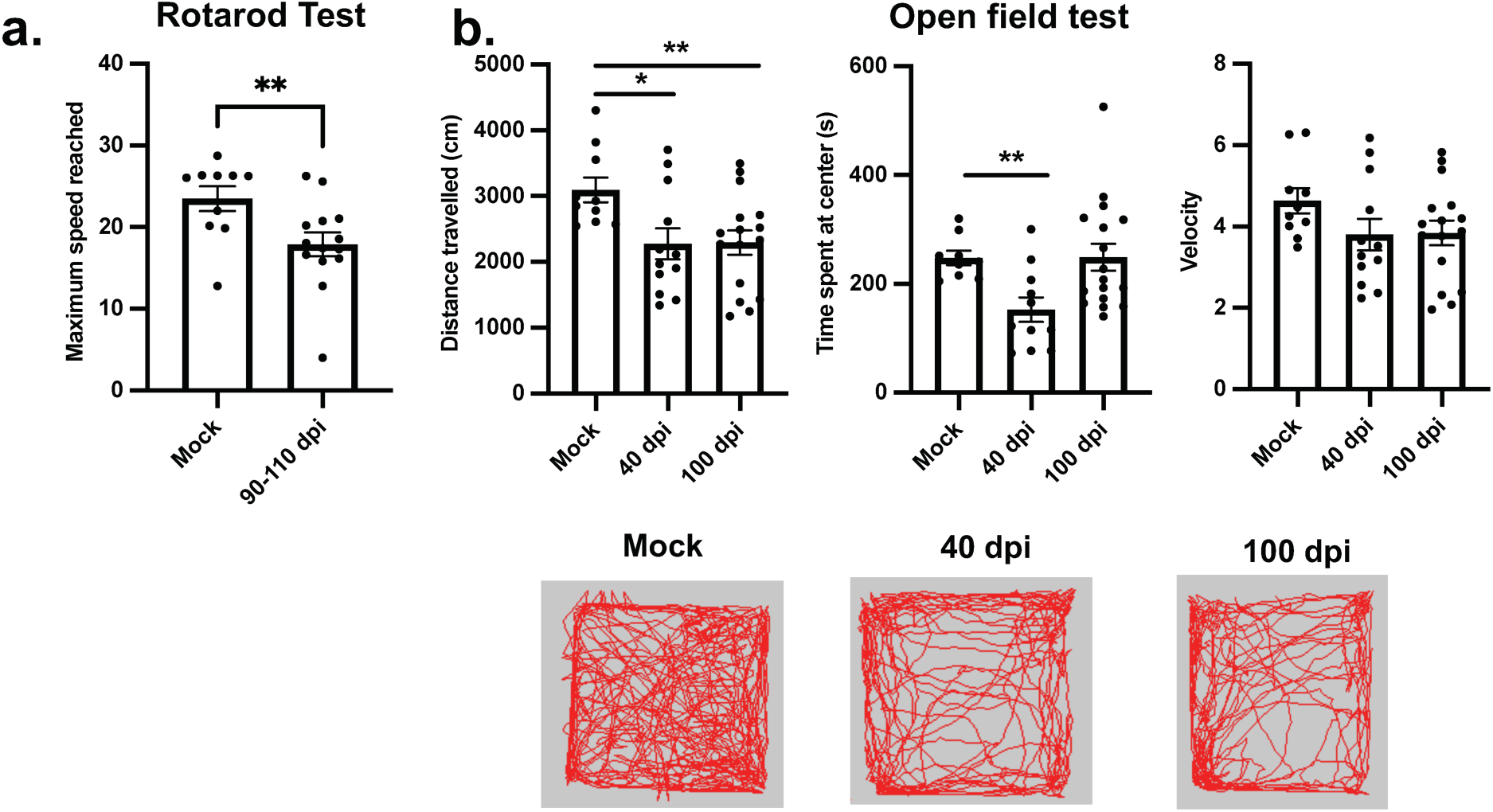
Behavioral manifestations. (a) Rotarod testing was performed in mock-infected and infected mice at 100 dpi. Mice were infected with 1000pfu SARS-COV-2. The highest speed sustained by mice without falling was recorded. Data represent mean ± SEM of results pooled from 2 independent experiments: mock (10 mice) and 90-110 dpi (12 mice). Data were analyzed using a Mann-Whitney U-test. \*\**P* < 0.01. (b) Open field testing of SARS-COV-2-mice was performed using mock-infected, 40dpi and 100 dpi mice. Data represent mean ± SEM of results pooled from 2 independent experiments: mock (10 mice), 40dpi (12 mice) and 120 dpi (16 mice). Data were analyzed using a Mann-Whitney U-test. \*\**P* < 0.01.

### Efficacy of nirmatrelvir and molnupiravir in combination in mitigating SARS-CoV-2-mediated disease in mice

While these data suggest that pathogenesis of neurological PASC is mediated at least in part by the host immune response, the precise role of SARS-CoV-2 is less clear. One possibility is that long term complications are dependent on the amount of the initial virus load, with the prediction that treatment with antiviral therapy would prevent or decrease disease. Nirmatrelvir and molnupiravir are FDA-approved for the treatment of SARS-CoV-2 at early times after infection. To examine whether antiviral drug treatment alleviated long-term term effects, we treated infected mice with nirmatrelvir (20mg/kg) and molnupiravir (20mg/kg) on a daily basis for 5 days beginning at day 0 of infection (Figure 6a). Drug treatment reduced SARS-CoV-2-induced weight loss and virus titers in the lungs at 2 dpi (Figure 6b-c). Levels of several pro-inflammatory cytokines and chemokines were also reduced by antiviral drug therapy in lungs (Extended data Figure 3). To assess if the observed reductions in virus titer and inflammation reversed behavioral changes in SARS-CoV-2-infected mice, we conducted open field testing on groups receiving either nirmatrelvir and molnupiravir or vehicle. We detected no significant improvement between the drug and vehicle-treated groups in the total distance traveled or the time spent in the center of the arena (Figure 6c). Additionally, we investigated TH expression in the mouse OB. Vehicle-treated mice exhibited reduced TH expression compared to mock-infected mice (Figure 6e). Mice treated with nirmatrelvir and molnupiravir showed no increase in TH expression compared to the vehicle-treated mice at 30 dpi (Figure 6e).

**Figure 6.**
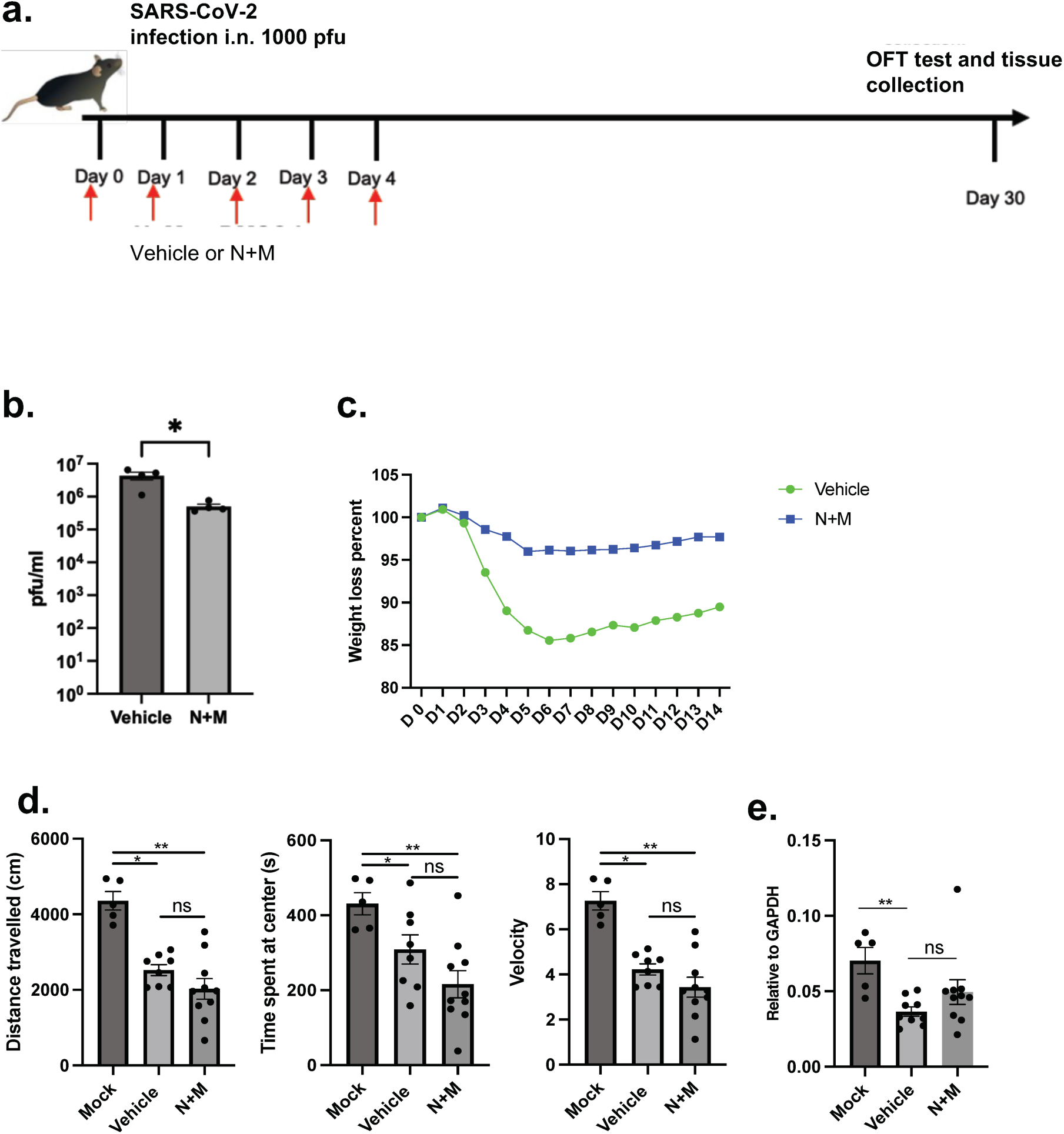
Nimratrelvir and molnupiravir reduce clinical manifestations and virus loads but do not reverse behavioral abnormalities. (a) Mice were infected with 1000pfu SARS-COV-2 and treated with nimratrelvir and molnupiravir (N+M) at the indicated times post infection. (b, c) Drug treatment reduced lung virus titers (b) and diminished weight loss (c). Data represent mean ± SEM of results pooled from 2 independent experiments. DMSO (n=9 mice), N+M (n=10 mice). (d) Open field testing was performed as described in Materials and Methods. Treatment with nimratrelvir and molnupiravir resulted in no improvement. Data represent mean ± SEM of results pooled from 2 independent experiments: mock (5 mice), DMSO (8 mice) and N+M (10 mice) (e) qPCR analysis shows TH mRNA levels in OB isolated from vehicle and drug-treated mice. Data represent mean ± SEM of results pooled from 2 independent experiments: mock (5 mice), DMSO (8 mice), and drug-treated (10 mice). Data were analyzed using a Mann-Whitney U-test. **P < 0.01.

### Substantia nigra from patients shows decreased TH+ cells

To determine the clinical relevance of our results showing decreased neurotransmitter expression in the murine SN, we obtained brain sections containing SN from deceased COVID-19 and uninfected control patients upon autopsy at times ranging from 4 to 56 days after SARS-CoV-2 infection. Demographics are provided in Table 1. There were no differences in the age or sex of COVID-19 versus uninfected control patients. Samples were stained for TH expression and numbers of TH+ cells were quantified. The data showed reduced numbers of TH+ cells in the SN of COVID-19 patients compared to non-infected control samples. In addition, in one patient analyzed at 12 day after diagnosis (patient #10), decreased pigmentation in the SN was noted, suggesting the loss of neuromelanin-positive neurons (Figure 7c). Next, we explored whether the reduced TH-staining resulted from preclinical Parkinson’s disease prior to infection. Given the gradual buildup of alpha-synuclein, we hypothesized that acute infection would unlikely cause its accumulation. To investigate this, we stained SN sections for alpha-synuclein. Most samples showed no staining, except for two with trace positive staining (Table 1). These data strongly suggest that SARS-CoV-2 infection was the cause of decreased TH-expression (Figure 7).

**Figure 7.**
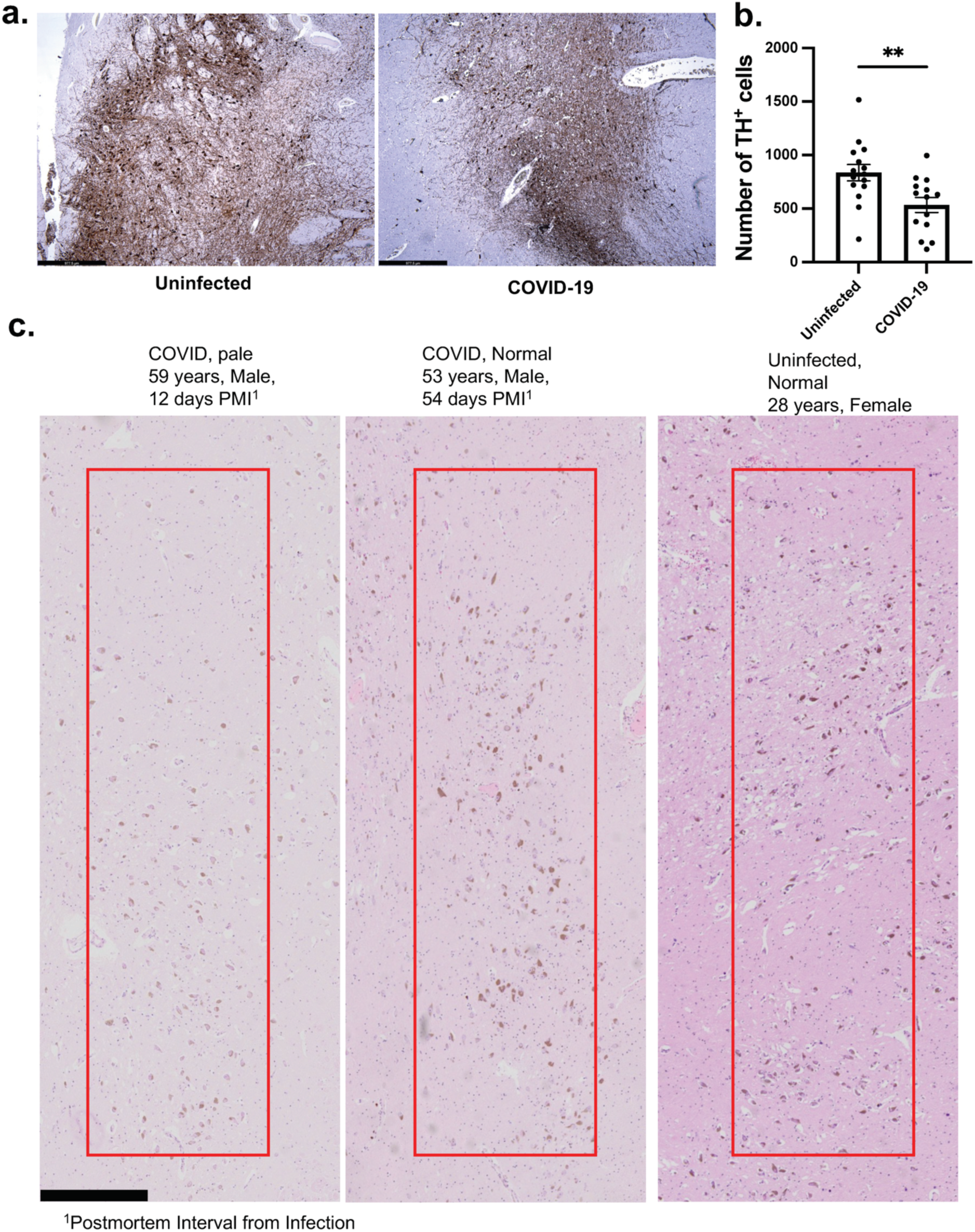
TH quantification in patients with COVID-19. (a) Immunostaining for TH was performed on the SN of autopsy samples from uninfected controls and patients with COVID-19 as described in Table 1 (b) Data show mean ± SEM. Uninfected control (n=15) and COVID-19 (n=14). Data were analyzed using a Mann-Whitney U-test. \*\**P* < 0.01. (c) H&E stained sections of the SN from COVID-19 patients were analyzed for changes in pigmentation. One COVID-19 patient had evident hypopigmentation (left), while pigmentation was normal in a second patient (middle) and an uninfected control (right). Boxed areas indicate sites of pigmentation.

**Table 1.** Patient Information.

| # | Age | Sex | | <sup>1</sup>PMI (days) | No. of TH+ cells | Pigmentation | α-syn staining |
| --- | --- | --- | --- | --- | --- | --- | --- |
| 1 | 64 | M | COVID-19 | 7 | 444 | Normal | Not Available |
| 2 | 73 | F | COVID-19 | 11 | 699 | Normal | Not Available |
| 3 | 61 | M | COVID-19 | 18 | 709 | Normal | Not Available |
| 4 | 65 | F | COVID-19 | 19 | 786 | Normal | Trace Positive |
| 5 | 80 | F | COVID-19 | 11 | 995 | Normal | Negative |
| 6 | 64 | M | COVID-19 | 4 | 188 | Normal | Negative |
| 7 | 67 | M | COVID-19 | 33 | 631 | Normal | Negative |
| 8 | 72 | M | COVID-19 | 30 | 351 | Normal | Trace positive <sup>2</sup>DRN |
| 9 | 53 | M | COVID-19 | 56 | 767 | Normal | Negative |
| 10 | 59 | M | COVID-19 | 12 | 379 | Reduced | Negative |
| 11. | 72 | M | COVID-19 | 10 | 675 | Not Available | Not Available |
| 12. | 73 | M | COVID-19 | 15 | 180 | Normal | Negative |
| 13. | 64 | F | COVID-19 | 23 | 120 | Not Available | Not Available |
| 14 | 86 | M | COVID-19 | 6 | 608 | Not Available | Not Available |
| 15 | 60 | F | Uninfected |  | 214 |  |  |
| 16 | 73 | F | Uninfected |  | 1053 |  |  |
| 17 | 62 | M | Uninfected |  | 1026 |  |  |
| 18 | 80 | F | Uninfected |  | 615 |  |  |
| 19 | 74 | M | Uninfected |  | 938 |  |  |
| 20 | 80 | F | Uninfected |  | 766 |  |  |
| 21 | 72 | M | Uninfected |  | 794 |  |  |
| 22 | 54 | M | Uninfected |  | 514 |  |  |
| 23 | 61 | F | Uninfected |  | 708 |  |  |
| 24 | 61 | M | Uninfected |  | 1517 |  |  |
| 25 | 69 | M | Uninfected |  | 821 |  |  |
| 26 | 67 | F | Uninfected |  | 721 |  |  |
| 27 | 76 | M | Uninfected |  | 878 |  |  |
| 28 | 75 | F | Uninfected |  | 1124 |  |  |

732 <sup>1</sup>Post mortem interval (interval between first positive PCR test and autopsy)733 <sup>2</sup>Dorsal raphe nucleus

## Discussion

The long-term consequences of COVID-19, PASC, continue to present significant challenges and are poorly understood^18,19,62^. Here we show that biochemical and behavioral changes persist in the mouse brain for several months after infection in the absence of infectious virus or viral RNA or protein. mRNA expression analyses of brain CD11b cells demonstrate evidence of prolonged inflammation, which coupled with changes in dopamine neuron levels, supports the hypothesis that the host immune response is a major if not the primary cause of the observed changes and by extension, PASC. We found vulnerability of dopaminergic cells in the OB and SN, two areas in the brain that are prominently affected in human neurodegenerative disease. Although changes in the OB during early stages of infection are not surprising as SARS-CoV-2 infects sustentacular cells in patients and experimentally infected animals^30–32,63^, the persistence of inflammatory responses in the OB, even after the clearance of virus, underscores the complex and enduring nature of PASC. Notably, the decrease in TH expression, indicative of diminished dopaminergic activity, and in hypopigmentation in the SN, observed in 1/11 patients, suggests a potential link to neurodegenerative processes^53–55,64^. Clinical studies support a role for the inflamed OB as a primary region associated with the development of neurodegenerative pathology^41–43^. Furthermore, quantitative analysis of the OB from COVID-19 patients revealed significantly decreased size, consistent with atrophy^65–67^. Additionally, infectious virus and viral RNA and protein cannot be detected in the olfactory mucosa at later times after infection^30^. A fraction of SARS-CoV-2-infected cells survive the acute infection in mice, but very few of these cells can be detected in the olfactory epithelium by 20 dpi^68^. Collectively, these results indicate that, whether or not neurological disease is triggered by sustentacular cell infection, continued infection or other abnormalities in the OE are not required for persisting dysfunction.

Activated microglia and neuroinflammation has been previously reported in SARS-CoV-2-infected patients and experimental animals including hamsters and macaques^38,60,69–72^. We show that microglial activation, a known marker of neuroinflammation, is observed in the OB and elsewhere in the SARS-CoV-2-infected brain. Consistent with our study, microglial activation was observed in infected hamsters in the olfactory nerve layer (ONL), glomerular and external plexiform layers (EPL) of the OB and persisted for as long as two weeks following infection^73^. We did not observe astrocytic hypertrophy, which was heightened in the SARS-CoV-2 infected hamster OB^73^, indicating that sustained microglial activation is sufficient to induce a protracted immune response and may play a role in the observed behavioral and biochemical alterations.

While diminished numbers of TH-immunoreactive neurons and increased pro-inflammatory molecule expression in the mouse OB at several months after infection was not expected, even more striking, were changes in the SN, a target region in human neurodegenerative disease. We detected heightened inflammation and a significant reduction in TH-positive neurons in conjunction with behavioral changes referable to the SN in the absence of viral infection. Decreases in TH-immunoreactive neurons in the SN signified loss of dopaminergic neurons, a hallmark of PD^54,56^. These findings, suggesting a link between the neuroinflammatory microenvironment and dopaminergic neuron loss, further support the notion that the host immune response following SARS-CoV-2 infection contributes to PD-like alterations.

Analyses of human bain samples also revealed decreased TH expression in the SN of deceased patients compared to controls indicating similar findings to those observed in mice (Figure 7). Our results are also in agreement with a recent report showing a reduction in neuromelanin-positive and TH-positive neurons in the SN in a cohort of deceased COVID-19 patients^74^. Previous studies highlighted the presence of neuroinflammation in the brains of COVID-19 patients, with CCL11 postulated to have a prominent negative effect on cognitive function^22^. In addition, elevated levels of IL-1b and IL-6 were detected in the brains of SARS-CoV-2-infected patients and hamsters and in hamsters were shown to contribute to impaired neurogenesis in the hippocampus^37^. In other studies, SARS-CoV-2 infection resulted in demyelination, disruptions in neurotransmitter synthesis, microgliosis, and an increase in alpha-synuclein levels^22,38,60,70–72,75^. A separate investigation into long COVID reported a decrease in cortical thickness in survivors of COVID-19, supporting a role for SARS-CoV-2 infection on the brain^35^.

The 1918 influenza pandemic led to a significant increase in cases of postencephalitic parkinsonism. This observation is consistent with our findings and suggest that basal ganglia dopaminergic (DA) neurons are especially susceptible to damage caused by influenza A virus or SARS-CoV-2 or associated immune responses^46–48,54,74^. Collectively, these results raise concerns that a long-term consequence of COVID-19 will be an increase in numbers of patients with neurodegenerative diseases. Notably, there are already several case reports of Parkinsonism following COVID-19, even though the pandemic began only 4 years ago^76,77^.

A key question is whether anti-viral treatment early in infection would decrease the amount of damage that is observed in the brain. While treatment with nirmatrelvir and molnupiravir in mice decreased virus titers and associated inflammatory responses (Figure 6), it was ineffective in alleviating neurological disease. Similarly, treatment with Paxlovid (nirmatrelvir/ritonavir) did not decrease the incidence of PASC in patients^78,79^, raising questions about the precise role of SARS-CoV-2 in the development of neurological sequelae. One possibility is that infection, even if transient, exacerbates sensitivity to environmental factors and accelerates neurodegeneration. Alternatively, it is also possible that the OB and SN are especially prone to persistent inflammation and therefore to higher levels of tissue damage and neurological disease.

In conclusion, our observations emphasize the intricate relationship between the role of virus infection, persistent inflammation, neurotransmitter dysregulation, and behavioral changes, in the development of neurological PASC. Furthermore, antiviral treatment reduced viral load and inflammation yet failed to prevent neurological dysfunction further pointing out the complex nature of the role of the virus. Minimizing the neuroinflammatory response, especially at early times after infection, may be critical for ameliorating neurological PASC.

## ONLINE CONTENTS

Any methods, Nature Research reporting summaries, source data, extended data, acknowledgments, peer review information; details of author contributions and competing interests; and statements of data and code availability are available at https://doi.org/.

## Material and Methods

### Animals and viruses

4-5 months old C57BL/6N mice (Charles River Laboratories) were used in all studies. Mice were infected intranasally with SARS-COV-2 (1 × 10^3^ pfu) as previously described^49^.

### Study approval

All animal studies were approved by the University of Iowa IACUC and met the stipulations of the Guide for the Care and Use of Laboratory Animals (National Academies Press, 2011). All human autopsy samples were obtained under an IRB-approved protocol (#202011287 (postmortem)) and after obtaining written informed consent from the participant or the family.

### Confocal imaging

For immunofluorescence assays, OB were fixed in zinc formalin and embedded in paraffin. Sections were deparaffinized and processed for citrate-based antigen retrieval (Vector Laboratories) according to the manufacturer’s protocol. Sections were washed 3 times for 5 minutes in PBS before treatment with 0.1% Triton X-100 in PBS for 20 minutes. Sections were then rinsed in PBS followed by incubation with CAS block (Invitrogen, Thermo Fisher Scientific) for 60 minutes. Primary antibodies against Iba1 (Wako, 1:1,000), and TH (Novus, 1:1,000) were used. Sections were rinsed before incubation with a 1:1,000 dilution of an appropriate Alexa Fluor 546–conjugated (catalog A11018) or A488-conjugated (catalog A11070) goat anti-mouse or anti-rabbit antibody (Thermo Fisher Scientific). After a final wash with PBS, slides were mounted with Vectashield antifade reagent containing DAPI (Vector Laboratories). Images were obtained using a Leica DM 4B fluorescence microscope. Three different areas were imaged from every brain section for cell counting. ImageJ (NIH) was used for image processing and cell enumeration.

Alpha-synuclein staining was done using clone EP1536Y (Abcam) at 1:100 using high pH antigen retrieval, with a 15-minute primary and polymer incubations on a Ventana auto-stainer instrument in the College of American Pathologists (CAP)-certified University of Iowa Diagnostic Histopathology Laboratory.

### RT-qPCR

Mice were deeply anesthetized with ketamine/xylazine and perfused transcardially with PBS. OB and SN were isolated and collected in Trizol (Thermo Fisher Scientific). mRNA expression levels were analyzed by quantitative PCR (qPCR). The primer sets used for PCR are listed in Table 2. SARS-CoV-2 N primers were purchased from IDT (catalog 10007032). Expression levels were normalized to GAPDH by the following CT equation: ΔCT = CT of the gene of interest – CT of GAPDH. All results are shown as a ratio to GAPDH calculated as 2^−ΔCt^.

**Table 2.**
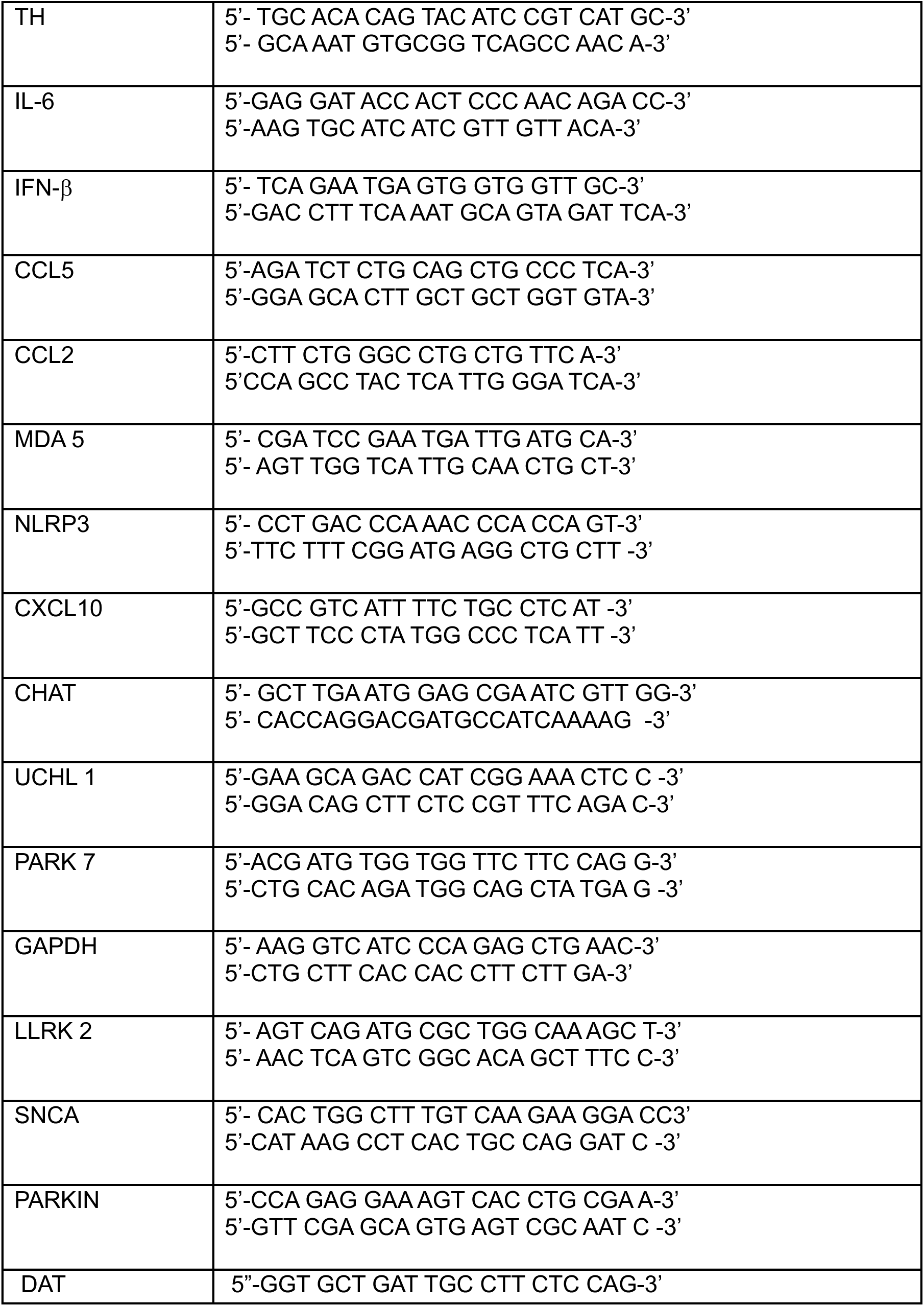
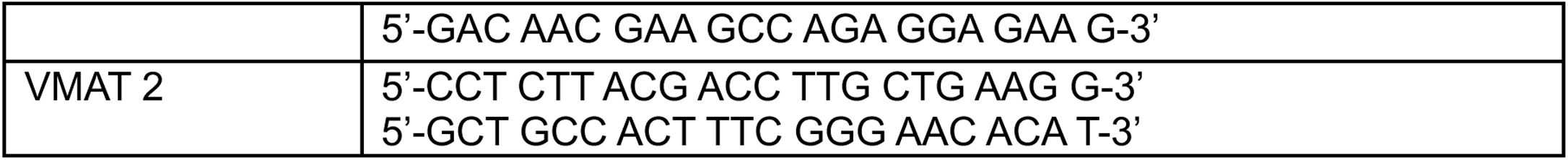
List of Primers.

### Gene expression profiling

CD11b+ cells from brains were sorted using magnetic beads (Miltenyi Biotech) and RNA was extracted using an RNeasy micro kit (Qiagen) per the manufacturer’s instructions. Four samples per group were analyzed. Subsequent library preparation and sequencing were performed at the University of Minnesota Genomics Center. RNA isolates were quantified using a fluorimetric RiboGreen assay, and RNA integrity was assessed using capillary electrophoresis (Agilent 2100 Bioanalyzer) to generate an RNA integrity number. Samples with RNA integrity number values >5.5 and at least 250 pg of total RNA were then used to generate 12 unique dual-indexed libraries using Takara/Clontech Stranded Total RNA-Seq Kit v2 -Pico Input Mammalian reagents. Briefly, between 250 pg and 10 ng of total RNA was fragmented and then reverse transcribed into cDNA using random primers, with a template-switching oligonucleotide incorporated during cDNA synthesis to allow for full-length cDNA synthesis and for retention of strand specificity. Illumina sequencing adapters and barcodes were then added to the cDNA by PCR, followed by cleavage of ribosomal cDNA. Uncleaved fragments were then enriched by PCR for 12–16 cycles. The final library size distribution was again validated using capillary electrophoresis and quantified using fluorimetry (PicoGreen). Indexed libraries were then normalized and pooled for sequencing sequencing on a NextSeq 2000.

2 x 50bp FastQ paired end reads for 8 samples (n=39.6 Million average reads per sample) were trimmed using Trimmomatic (v 0.33) enabled with the optional “-q” option; 3bp sliding-window trimming from 3’ end requiring minimum Q3Quality control on raw sequence data for each sample were performed with FastQC. Read mapping was performed via Hisat2 (v2.1.0) using the mouse genome (GRCm39 v109) as reference. Gene quantification was done via Feature Counts for raw read counts. Differentially expressed genes were identified using the edgeR (negative binomial) feature in CLCGWB (Qiagen, Redwood City, CA) using raw read counts. The list of Differentially Expressed Genes (DEGs) was generated based on a minimum 2x Absolute Fold Change and FDR corrected p < 0.05. Differentially expressed genes were identified using the edgeR (negative binomial) feature in CLCGWB (Qiagen) using raw read counts. Heatmaps using designated sets of differentially expressed genes were generated using pheatmap (R). Ingenuity Pathway Analysis software (Qiagen) was used to analyze biological pathways.

### Mouse behavior

For rotarod testing, mice were habituated with at 8rpm for several trials until they adapted to the rod for two consecutive days. After habituation, mice were analyzed at accelerating speed for three trials per mouse. The highest speed reached by mice before falling was recorded. For open field testing, the open field arena (40 cm X 40cm X 20 cm height, white background) was placed in a room with an indirect artificial light source. The arena was cleaned thoroughly with a 5% alcohol/water solution between each mouse to minimize odor cues. Mice from different groups were tested randomly throughout the trials so that mice from a single group were not tested consecutively. One mouse at a time was placed in the center of the arena and spontaneous behavior was recorded for 10 min. *V*ideos were evaluated using an Ethovision XT video tracking system (Noldus Information Technology, the Netherlands) to measure the distance moved (cm), average speed (cm/s), time moving (s), and time spent in the periphery/center of the arena (s)

### Treatment with antiviral therapy

Nirmatrelvir and molnupiravir were purchased from MedChemExpress (Monmouth Junction, New Jersey, USA). Stock solutions were prepared in DMSO. 20mg/kg of each compound was injected intraperitoneally into the mice once daily for five days starting at 0 dpi.

### Statistical analysis

Data were analyzed using a Mann-Whitney U-test. P < 0.05 was considered significant. Data in graphs are presented as mean ± SEM.

### Data availability

Complete RNA-seq data were deposited in the National Center for Biotechnology Information’s Gene Expression Omnibus database under number GSE254984 https://www.ncbi.nlm.nih.gov/geo/query/acc.cgi?acc=GSE254984

## Author’s contributions

The study was designed by SP and AKV. Experiments were conducted by AKV, SL, ED, LCL. AKV, MH, QQ, CRY, MWA and SP acquired and analyzed data. JE helped with RNAseq data analysis. LCL, MH provided reagents. Manuscript was initially prepared by AKV and SP. All of the authors revised and approved the final manuscript.

## Acknowledgments

We thank Mariah R Leidinger (Comparative Pathology Laboratory, University of Iowa), Shane Heiney (Neural Circuit Behavior Core, University of Iowa) and Kurt Bedell for technical support. Supported in part by grants from the NIH (R01 NS36592, P01 AI060699, R01 AI129269 awarded to SP; RF1 AG078297, awarded to MWA). Biobanking in the Iowa NeuroBank Core is supported in the Iowa Neuroscience Institute at the University of Iowa Roy J. and Lucille A Carver College of Medicine by the Roy J. Carver Charitable Trust.

## Extended Data Figure legend

**Extended data Figure 1.**
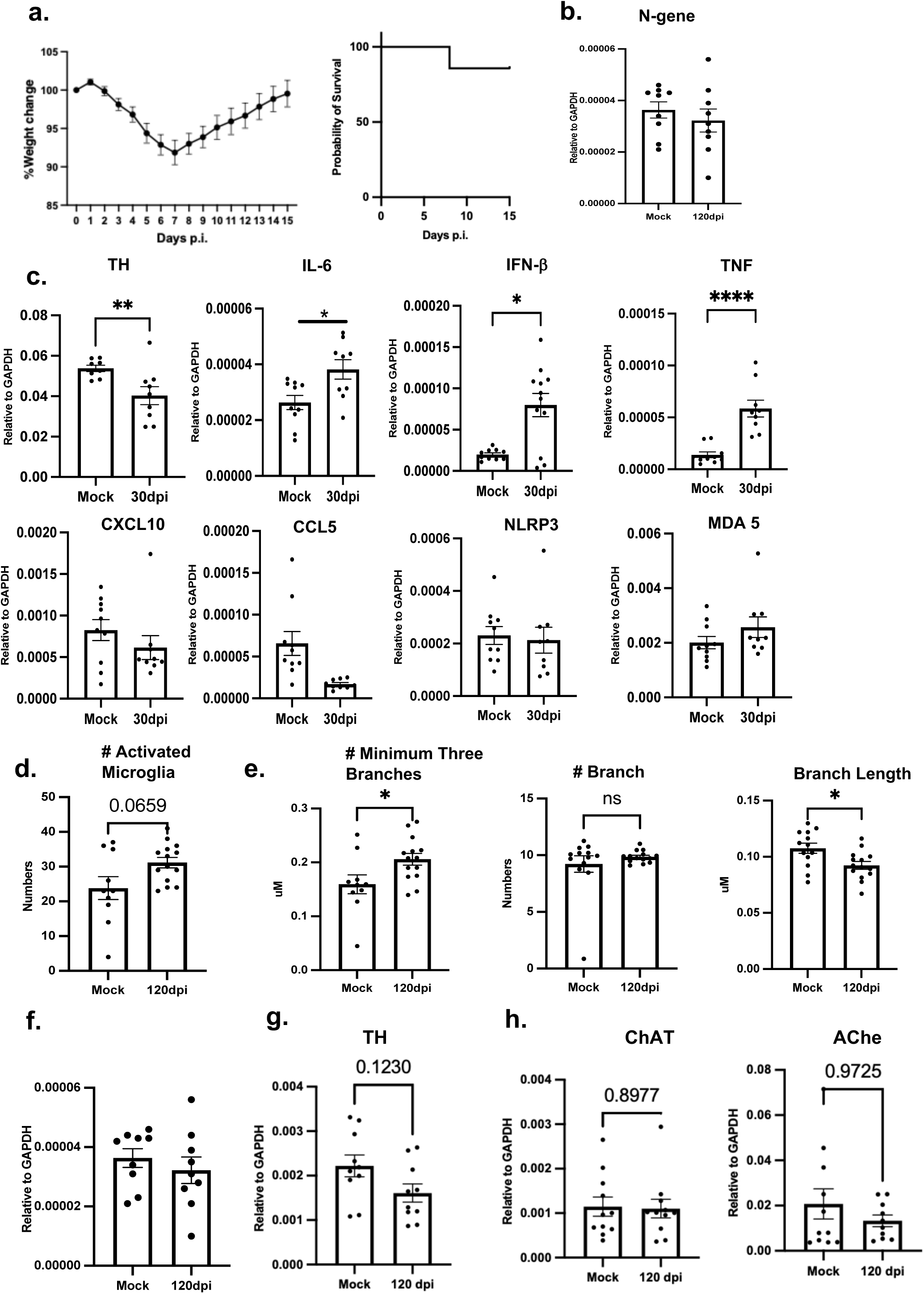
Clinical manifestations, microglia activation and neurotransmitter expression in SARS-CoV-2 infected mice. (a) Weight loss and survival in mice infected with 1000pfu of SARS-COV-2. (b) N and GAPDH mRNA levels in the OB were determined by qPCR at 120 dpi. Data represent mean ± SEM of results pooled from 2 independent experiments; mock (9 mice) and 120 dpi (9 mice) and were analyzed using a Mann-Whitney U-test. (c) OB mRNA was analyzed for TH expression by qPCR. Data represent mean ± SEM of results pooled from 2 independent experiments: mock (8-9 mice) and 30 dpi (9-11 mice). Data were analyzed using a Mann-Whitney U-test, *P < 0.05. (d) Microglia were quantified and analyzed for phenotypic changes consistent with activation as described in Materials and Methods. Data were analyzed using a Mann-Whitney U-test. *P < 0.05. (e) Summary data of microglia skeleton changes. Data were analyzed using a Mann-Whitney U-test. *P < 0.05. (f) N and GAPDH mRNA levels in the SN were determined by qPCR at 120 dpi. Data represent mean ± SEM of results pooled from 2 independent experiments; mock (9 mice) and 120 dpi (9 mice) and were analyzed using a Mann-Whitney U-test. (g-h) TH, ChAT and AChE mRNA expression in brains from which the OB, substantia nigra and cerebellum were removed. Data represent mean ± SEM of results pooled from 2 independent experiments: mock, 120 dpi (5 mice). Data were analyzed using a Mann-Whitney U-test.

**Extended Data Figure 2.**
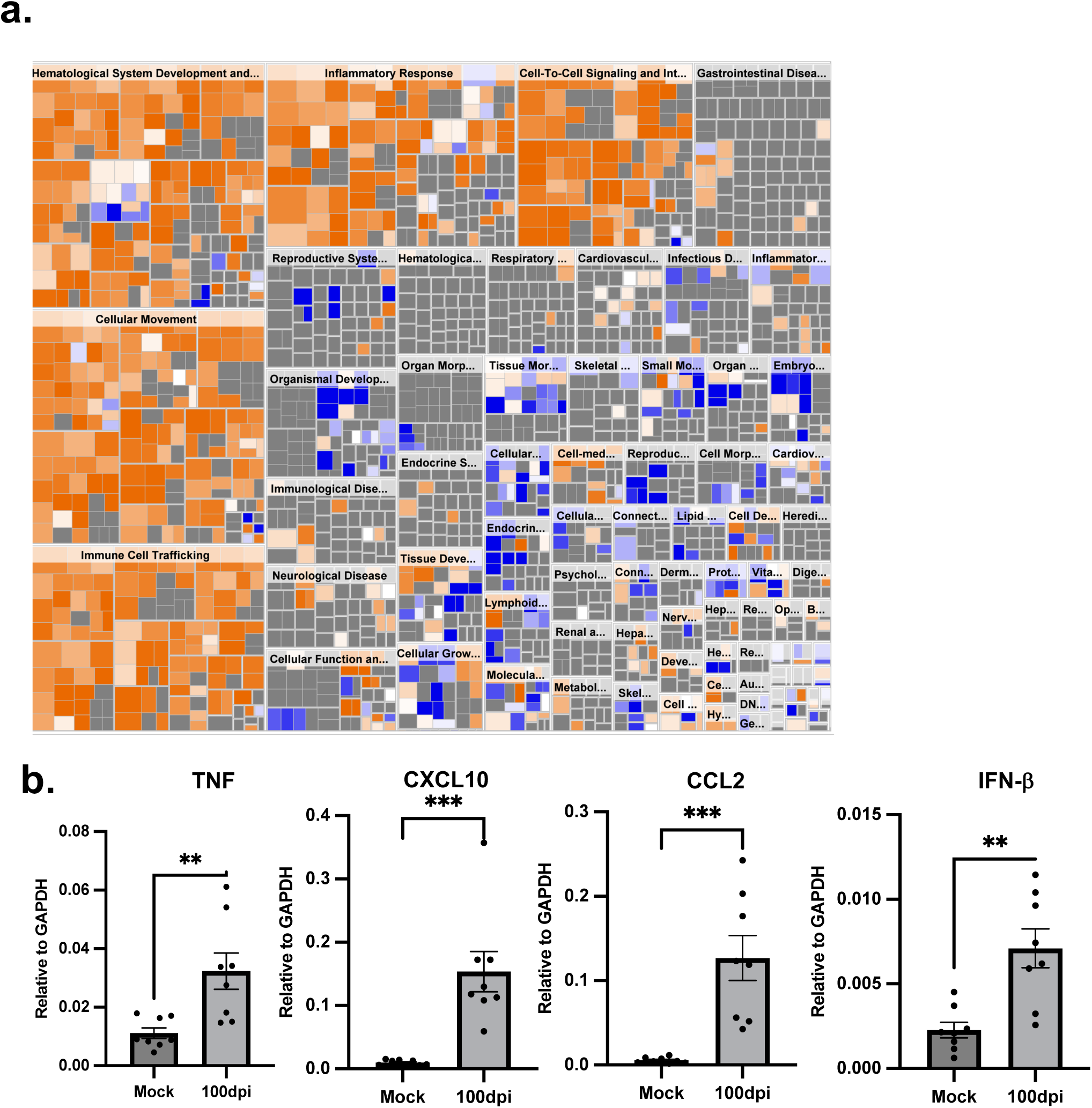
Differential gene expression in CD11b+ cells from SARS-COV-2 and mock-infected brains. CD11b+ cells were prepared from the brains of uninfected and infected mice at 100 dpi. RNA was prepared and analyzed by RNAseq as described in Materials and Methods. (a) Disease-specific canonical pathways upregulated in CD11b+ cells isolated from infected mouse brains were identified by IPA. Upregulation and downregulation of disease and functions are represented by blue and red color respectively. Inflammatory pathways and cellular trafficking pathways were upregulated. The ratio of differentially expressed genes for disease and functions is shown with Benjamini-Hochberg corrected p-values < 0.05, |fold-change| ≥ 2.5. (b) mRNA levels of specific pro-inflammatory genes are shown. Data represent mean ± SEM of results pooled from 2 independent experiments: mock (8 mice), 100 dpi (8 mice). Data were analyzed using a Mann-Whitney U-test. **P < 0.01, ***P < 0.001

**Extended Data Figure 3.**
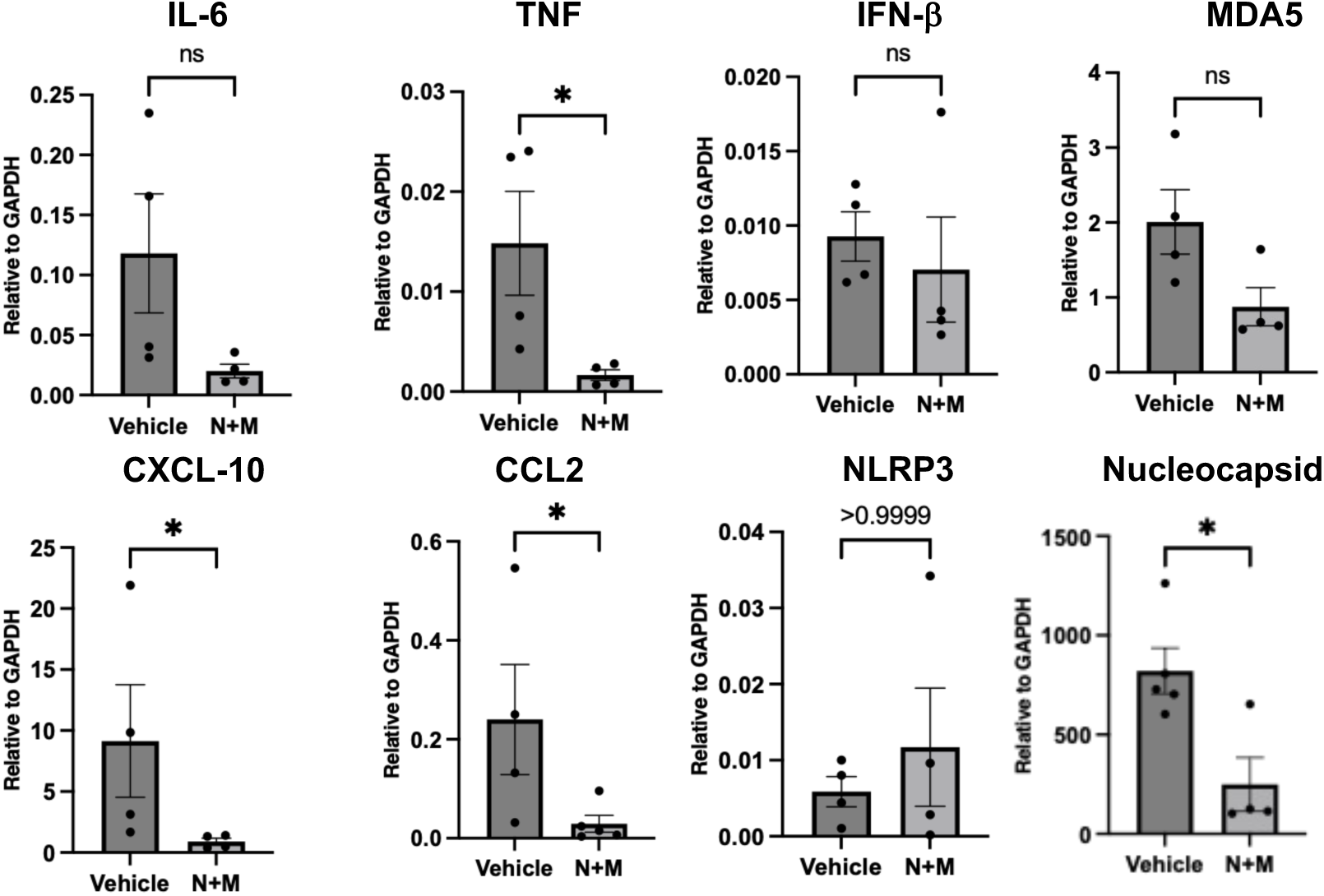
Nimratrelvir and molnupiravir treatment reduces mRNA levels of pro-inflammatory genes and N gene. Levels of mRNA were analyzed using qPCR at day 2 dpi. Data represent mean ± SEM of results pooled from 5 mice and were analyzed using a Mann-Whitney U-test. **P < 0.01

